# A rationally designed c-di-AMP FRET biosensor to monitor nucleotide dynamics

**DOI:** 10.1101/2021.02.10.430713

**Authors:** Alex J. Pollock, Philip H. Choi, Shivam A. Zaver, Liang Tong, Joshua J. Woodward

## Abstract

3’3’-cyclic di-adenosine monophosphate (c-di-AMP) is an important nucleotide second messenger found throughout the bacterial domain of life. C-di-AMP is essential in many bacteria and regulates a diverse array of effector proteins controlling pathogenesis, cell wall homeostasis, osmoregulation, and central metabolism. Despite the ubiquity and importance of c-di-AMP, methods to detect this signaling molecule are limited, particularly at single cell resolution. In this work, crystallization of the *Listeria monocytogenes* c-di-AMP effector protein Lmo0553 enabled structure guided design of a Förster resonance energy transfer (FRET) based biosensor, which we have named CDA5. CDA5 is a fully genetically encodable, specific, and reversible biosensor which allows for the detection of c-di-AMP dynamics both *in vitro* and within live single cells in a nondestructive manner. Our initial studies identify a unimodal distribution of c-di-AMP in *Bacillus subtilis* which decreases rapidly when cells are grown in diluted Luria Broth. Furthermore, we find that *B. subtilis* mutants lacking either a c-di-AMP phosphodiesterase or cyclase have respectively higher and lower FRET responses, again in a unimodal manner. These findings provide novel insight into c-di-AMP distribution within bacterial populations and establish CDA5 as a powerful platform for characterizing new aspects of c-di-AMP regulation.

**Importance:** C-di-AMP is an important nucleotide second messenger for which detection methods are severely limited. In this work we engineer and implement a c-di-AMP specific FRET biosensor to remedy this dearth. We present this biosensor, CDA5, as a versatile tool to investigate previously intractable facets of c-di-AMP biology.

## INTRODUCTION

Bacterial growth, reproduction, and survival demand rapid, accurate, and coordinated responses to internal and environmental cues. These responses are commonly relayed by nucleotide second messengers which dynamically change concentration through rapid synthesis and degradation[^1–5^]. The nucleotide second messenger 3’3’-cyclic di-adenosine monophosphate (c-di-AMP) is unique in that it is essential in diverse bacterial genera and regulates clinically and industrially relevant processes such as osmotic stress responses, cell wall metabolism, and metabolic homeostasis[^1,6–26^]. C-di-AMP is known to be produced by five classes of di-adenylate cyclases, although most bacteria contain only one of two main cyclases: the membrane associated CdaA and DNA binding DisA[^1,9,10^]. *Bacillus subtilis*, the model organism used in this study, interestingly encodes *cdaA*, *disA*, and a spore restricted cyclase, *cdaS*. Due to redundancy, it is possible to delete individual cyclases in *B. subtilis* without dramatic phenotypic consequences. C-di-AMP is degraded by 4 classes of phosphodiesterase although most bacteria only encode one or two of these enzymes[^1,4,9,10^]. *Bacillus subtilis* encodes both *gdpP* and *pgpH*, which again, due to redundancy, allows for deletion of single phosphodiesterases without significant physiological defects.

Despite being a second messenger of growing interest, only a small handful of studies have explored the internal and external signals which regulate the activity of c-di-AMP cyclases and phosphodiesterases. Multi-hour exposure to glutamine, potassium, and light have been reported to impact signal levels but the mechanisms by which nucleotide levels change have either not been defined or have been linked to altered phosphodiesterase expression[^6,12,14,19,24–26^]. Additionally, investigations of protein-protein interactions influencing intracellular c-di-AMP levels have been performed elucidating DacA regulation by the cistronic GlmM and YbbR proteins[^27 28 29^]. To our knowledge, the only studies which clearly identified post-translational c-di-AMP responses to environmental conditions have been: first, that a wide array of bacteria when non-growing but energized quickly accumulate more c-di-AMP in low osmotic conditions relative to high osmotic conditions, and second, that (p)ppGpp, which accumulates during amino acid starvation, inhibits c-di-AMP phosphodiesterases[^14,16,17^]. Such foundational studies remain largely elusive because detection methods are underdeveloped.

C-di-AMP is currently quantified using either mass spectrometry or an enzyme immunoassay which are both relatively low throughput and expensive methods providing only a snapshot of the population average at a single time point[^30,31^]. Additionally, an RNA-aptamer based biosensor has also been developed to detect c-di-AMP[^32^]. Unfortunately, this biosensor has not been able to be utilized because it has a long k_off_ which obscures c-di-AMP decreases, suffers from stochasticity in fluorescent ligand uptake, and lacks an internal control. To partially remedy this dearth, we recently developed the CDA-Luc assay which is an inexpensive and higher throughput method for quantifying c-di-AMP[^33^]. Although we anticipate this method being invaluable to many investigations, it is still limited to destructive snapshots of the population average.

Thus, an intramolecular FRET biosensor for c-di-AMP is an ideal tool to complement existing techniques as it would allow for nondestructive and rapid resampling of c-di-AMP in single cells. Intramolecular FRET biosensors have been central to diverse investigations of nucleotide second messenger regulation in both eukaryotic and bacterial cells[^34–46^]. Most similarly to c-di-AMP, FRET biosensors have been used extensively to study the bacterial nucleotide second messenger 3’3’-cyclic di-guanosine monophosphate (c-di-GMP)[^34–43^]. These investigations have interrogated important phenomena which are currently intractable using current c-di-AMP detection methods including: sub-population responses to environmental conditions, mother/daughter cell heterogeneity, and regulation during infection.

Intramolecular FRET biosensors are fusion proteins which combine compatible fluorophores with a native binding protein for the ligand of interest[^47–53^]. The locations of these fluorophores are engineered such that ligand binding induces a conformational change which moves the donor fluorophore either closer to or further from the acceptor fluorophore generating altered energy transfer. This shift in energy transfer causes a change in the fluorescent signal quantified as the ratio of: energy transferred to and released by the acceptor fluorophore divided by energy released by the donor fluorophore. The FRET ratio reports on the binding state of the biosensor allowing for back calculation of the free ligand concentration within the solution or cell. Thus, FRET biosensors are powerful, entirely genetically encodable, internally controlled, and, due to their native effector protein scaffold, have physiologically relevant binding parameters.

In this work, we engineer and utilize CDA5: a FRET biosensor designed around the *Listeria monocytogenes* c-di-AMP binding protein, Lmo0553[^13^]. We demonstrate that CDA5 retains relevant native binding characteristics, produces a robust FRET response upon the addition of c-di-AMP, and successfully reports on the concentration of c-di-AMP in individual bacterial cells. Furthermore, we report that *B. subtilis* grown in diluted LB media rapidly decreases its intracellular c-di-AMP concentration and that c-di-AMP phosphodiesterase and cyclase mutants, as expected, have respectively higher and lower corresponding FRET signals. Additionally, we report that, in all conditions tested and unlike c-di-GMP, c-di-AMP concentrations in single cells follow a unimodal distribution. Our investigations not only identify a new facet of c-di-AMP biology but also establish CDA5 as a versatile platform which will facilitate a wealth of basic and applied investigations of the essential bacterial signaling molecule c-di-AMP.

## RESULTS

### Overall structure of Lmo0553 in complex with cyclic-di-AMP

We previously identified Lmo0553 as a *Listeria monocytogenes* protein of unknown function which binds c-di-AMP at physiologically relevant concentrations. Despite numerous attempts to investigate the function of Lmo0553, its physiology has remained elusive. Thus, we sought to determine its basis of recognition and molecular response to c-di-AMP. The crystal structure of Lmo0553 in complex with c-di-AMP was determined at 1.6 Å resolution, as well as the structure of free Lmo0553 at 2.48 Å resolution. The atomic models have good agreement with the crystallographic data and the expected bond lengths, bond angles, and other geometric parameters (Table 1).

**Table 1:**
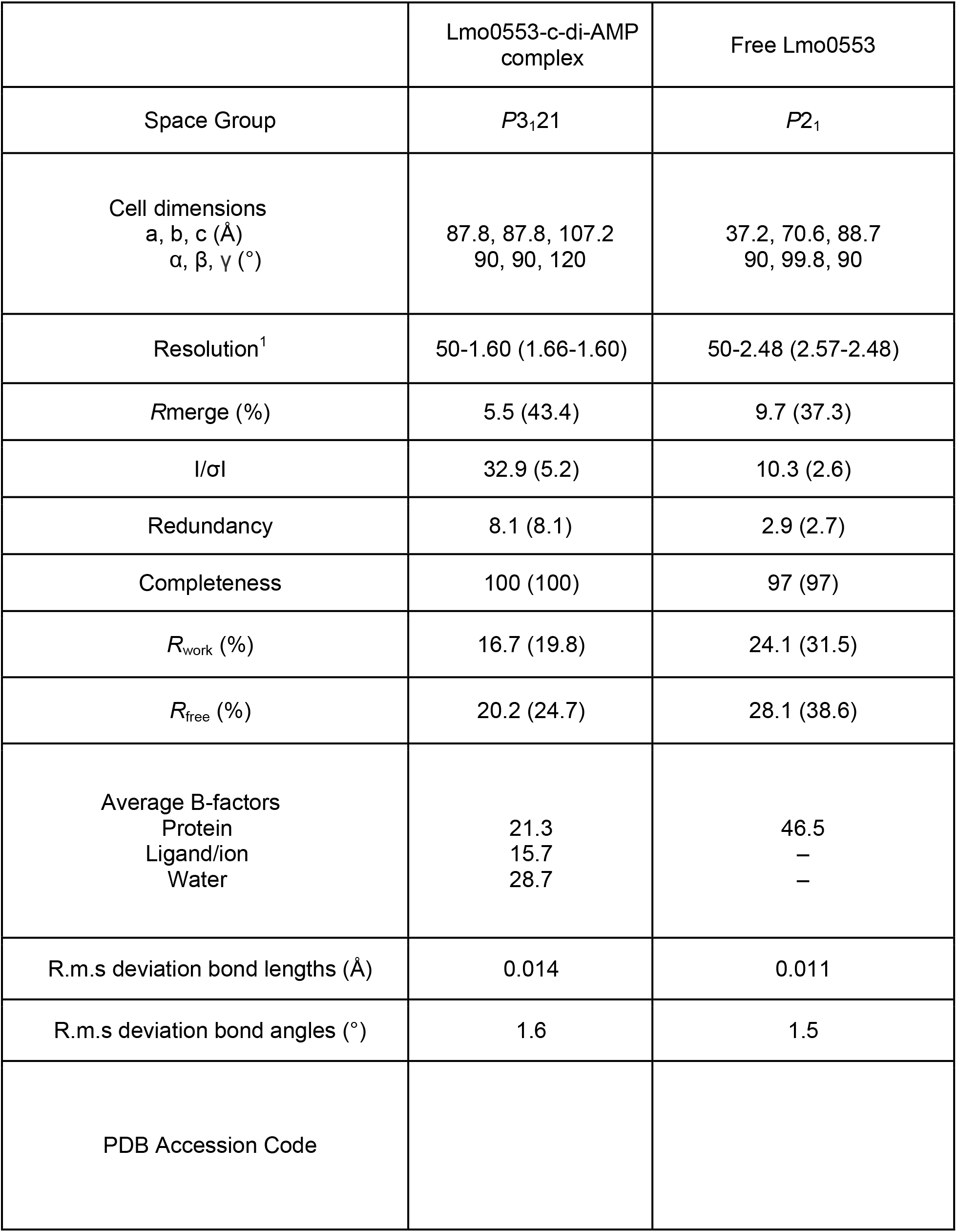
Summary of Crystallographic Information.

|  | Lmo0553-c-di-AMP<br>complex | Free Lmo0553 |
| --- | --- | --- |
| Space Group | $P3_121$ | $P2_1$ |
| Cell dimensions<br>a, b, c (Å)<br>$\alpha$, $\beta$, $\gamma$ (°) | 87.8, 87.8, 107.2<br>90, 90, 120 | 37.2, 70.6, 88.7<br>90, 99.8, 90 |
| Resolution<sup>1</sup> | 50-1.60 (1.66-1.60) | 50-2.48 (2.57-2.48) |
| Rmerge (%) | 5.5 (43.4) | 9.7 (37.3) |
| I/σI | 32.9 (5.2) | 10.3 (2.6) |
| Redundancy | 8.1 (8.1) | 2.9 (2.7) |
| Completeness | 100 (100) | 97 (97) |
| R<sub>work</sub> (%) | 16.7 (19.8) | 24.1 (31.5) |
| R<sub>free</sub> (%) | 20.2 (24.7) | 28.1 (38.6) |
| Average B-factors<br>Protein<br>Ligand/ion<br>Water | 21.3<br>15.7<br>28.7 | 46.5<br>—<br>— |
| R.m.s deviation bond lengths (Å) | 0.014 | 0.011 |
| R.m.s deviation bond angles (°) | 1.6 | 1.5 |
| PDB Accession Code |  |  |

**Table 2:**
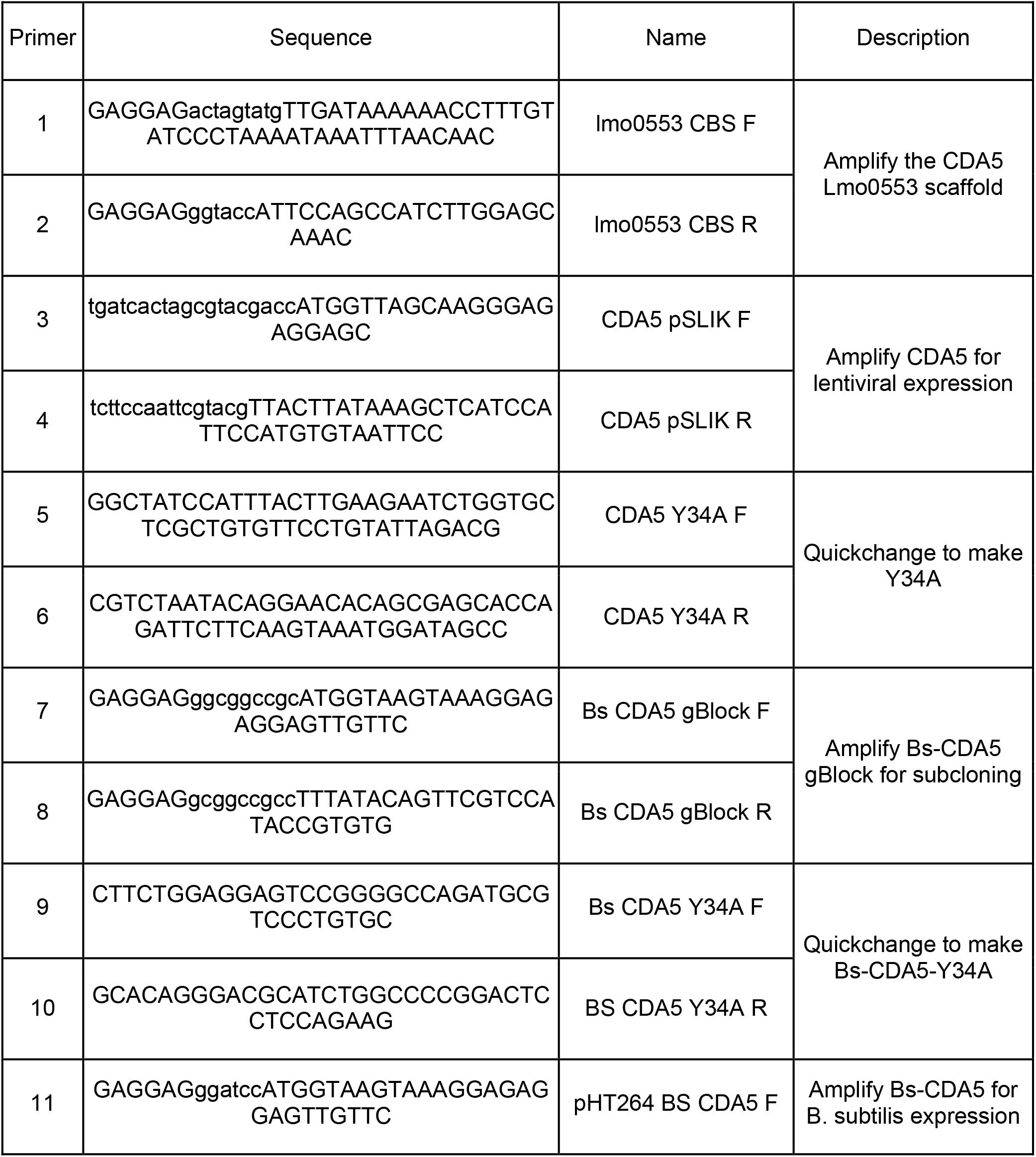
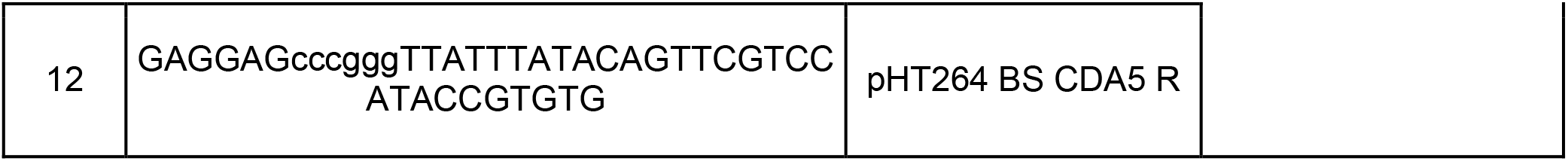
Primers.

**Table 3:**
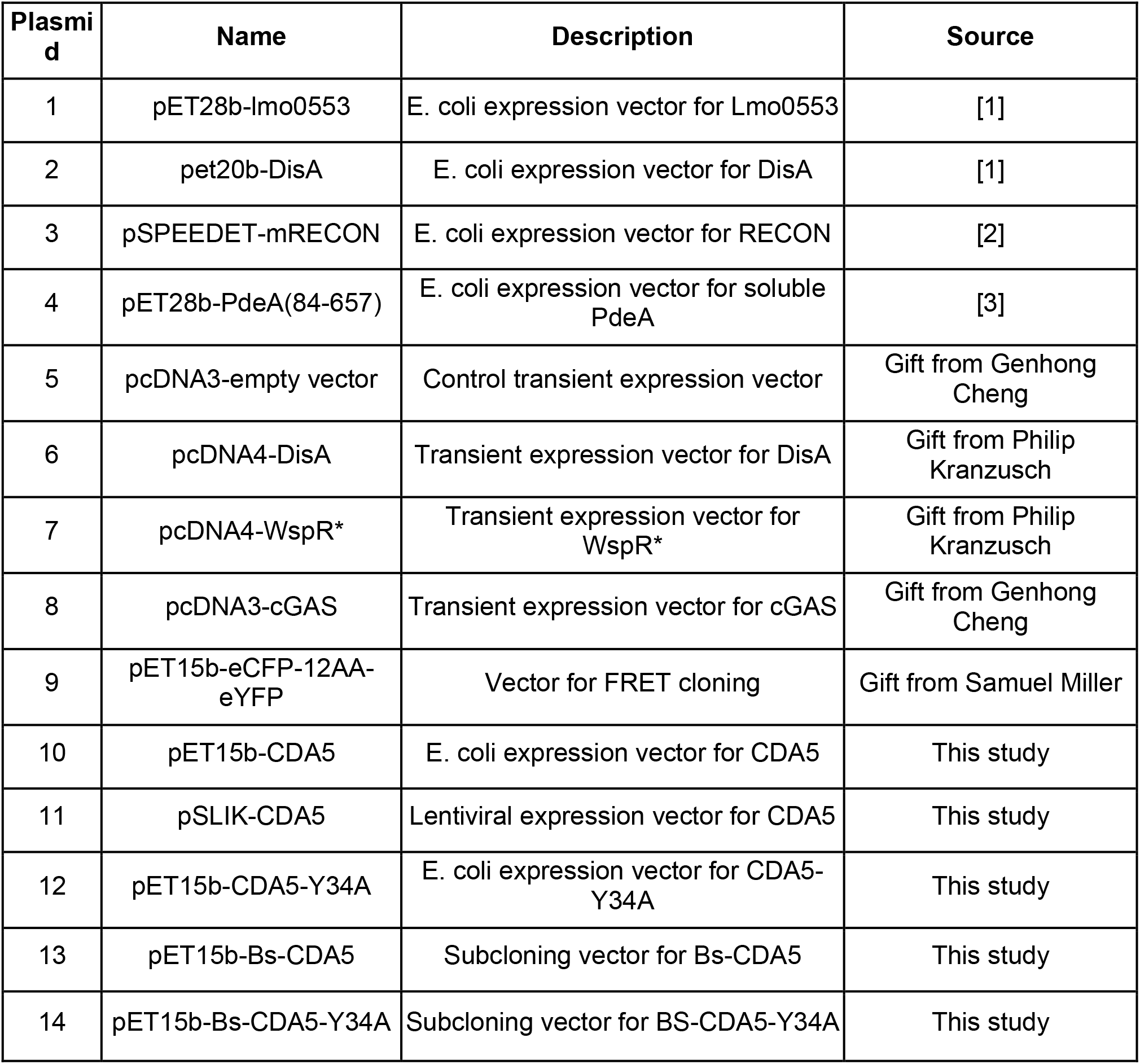
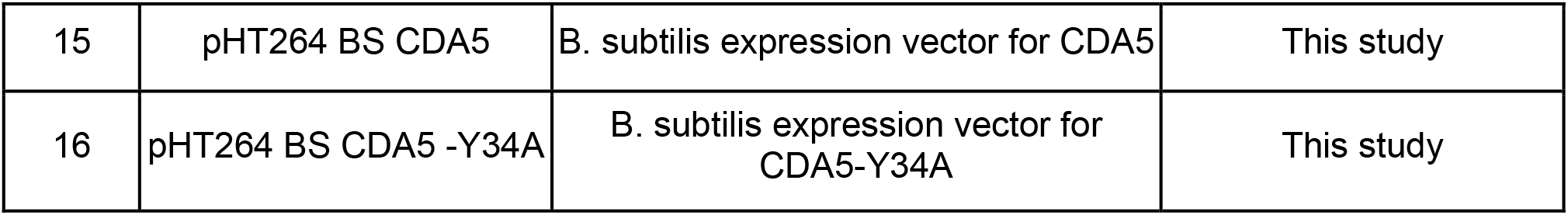
Plasmids. **Table 3 references:** 1) Sureka, K. *et al.* The cyclic dinucleotide c-di-AMP is an allosteric regulator of metabolic enzyme function. *Cell* **158**, 1389–1401 (2014). 2) McFarland, A. P. *et al.* Sensing of Bacterial Cyclic Dinucleotides by the Oxidoreductase RECON Promotes NF-κB Activation and Shapes a Proinflammatory Antibacterial State. *Immunity* **46**, 433–445 (2017). 3) Witte, C. E. *et al.* Cyclic di-AMP is critical for Listeria monocytogenes growth, cell wall homeostasis, and establishment of infection. *MBio* **4**, e00282–13 (2013).

**Table 4:**
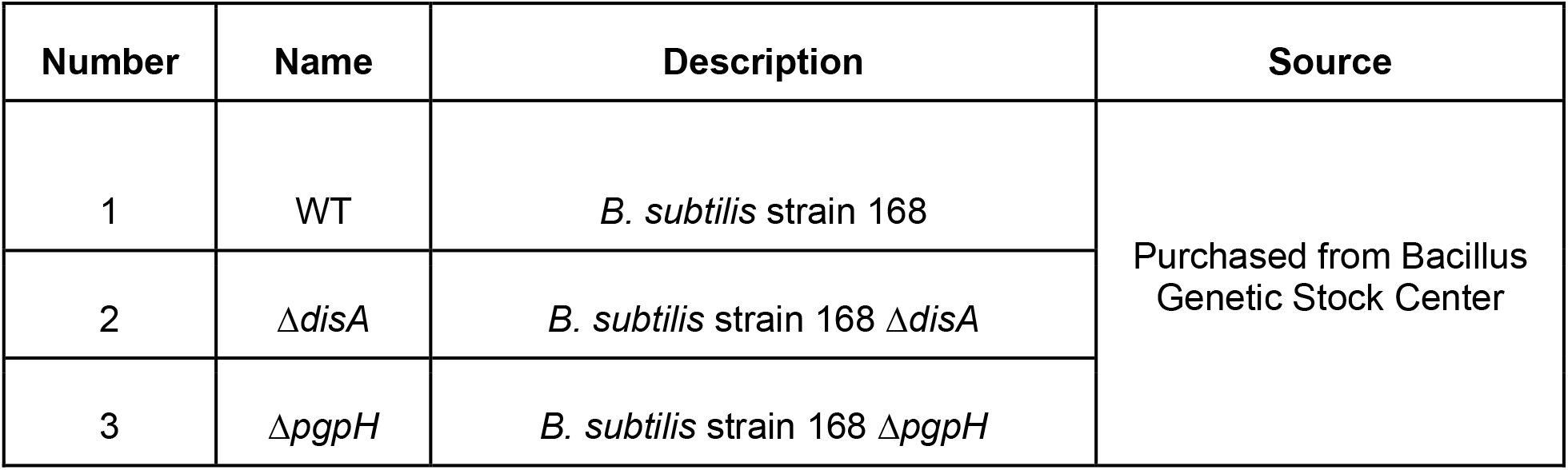
Strains.

| Number | Name | Description | Source |
| --- | --- | --- | --- |
| 1 | WT | *B. subtilis* strain 168 | Purchased from Bacillus Genetic Stock Center |
| 2 | $\Delta disA$ | *B. subtilis* strain 168 $\Delta disA$ | |
| 3 | $\Delta pgpH$ | *B. subtilis* strain 168 $\Delta pgpH$ | |

Lmo0553 contains tandem CBS motifs (Bateman domain) followed by an ACT domain. We observed a dimer of Lmo0553 in the crystals in which the Bateman domains of the two monomers contact each other in a head-to-head fashion, forming a disc-like dimer (Figs. 1A, 1B). This dimer is similar to other structures of CBS motif-containing proteins. As predicted, the C-terminal ACT domain of Lmo0553 adopts a ferredoxin fold (βɑββɑβ). This domain also forms a dimer, producing a structure consisting of an 8-stranded β-sheet packed against four ɑ-helices on one face. The Bateman domain dimer sits atop the β-sheet of the ACT dimer, forming an extensive interface. The Bateman and ACT domains are diagonally swapped in the dimer, such that the Bateman domain of monomer 1 packs against the ACT domain of monomer 2 (Fig. 1A). Two c-di-AMP molecules are bound in the central cavity of the Bateman domain dimer, one on each face of the disc (Fig. 1B). The dinucleotide adopts a folded conformation, with both bases in the *anti* configuration. The adenine of the first nucleotide (labeled 1 in Fig. 1C) is buried between the CBS1 and CBS2 motifs of a monomer. This adenine makes extensive interactions with the protein and is recognized specifically, with a hydrogen-bond to its N1 and N6 atoms. Tyr34 is p-stacked against one face of the adenine ring, while several hydrophobic residues flank the other face. In contrast, the adenine of the second nucleotide (labeled 2 in Fig. 1C) makes few direct interactions with the protein. The most notable interaction of the second nucleotide is the 5-prime phosphate which interacts with the side chain of Arg35. In the free as compared to c-di-AMP bound Lmo0553 structure, the side chain of Arg35 assumes a different conformation and occupies the binding site of c-di-AMP (Fig. 1D). In the c-di-AMP complex structure, residues that contact the adenine base of the first nucleotide show substantial conformational differences as well. These changes in the binding site are likely the trigger for the overall changes in the organization of the dimer upon c-di-AMP binding.

**Figure 1:**
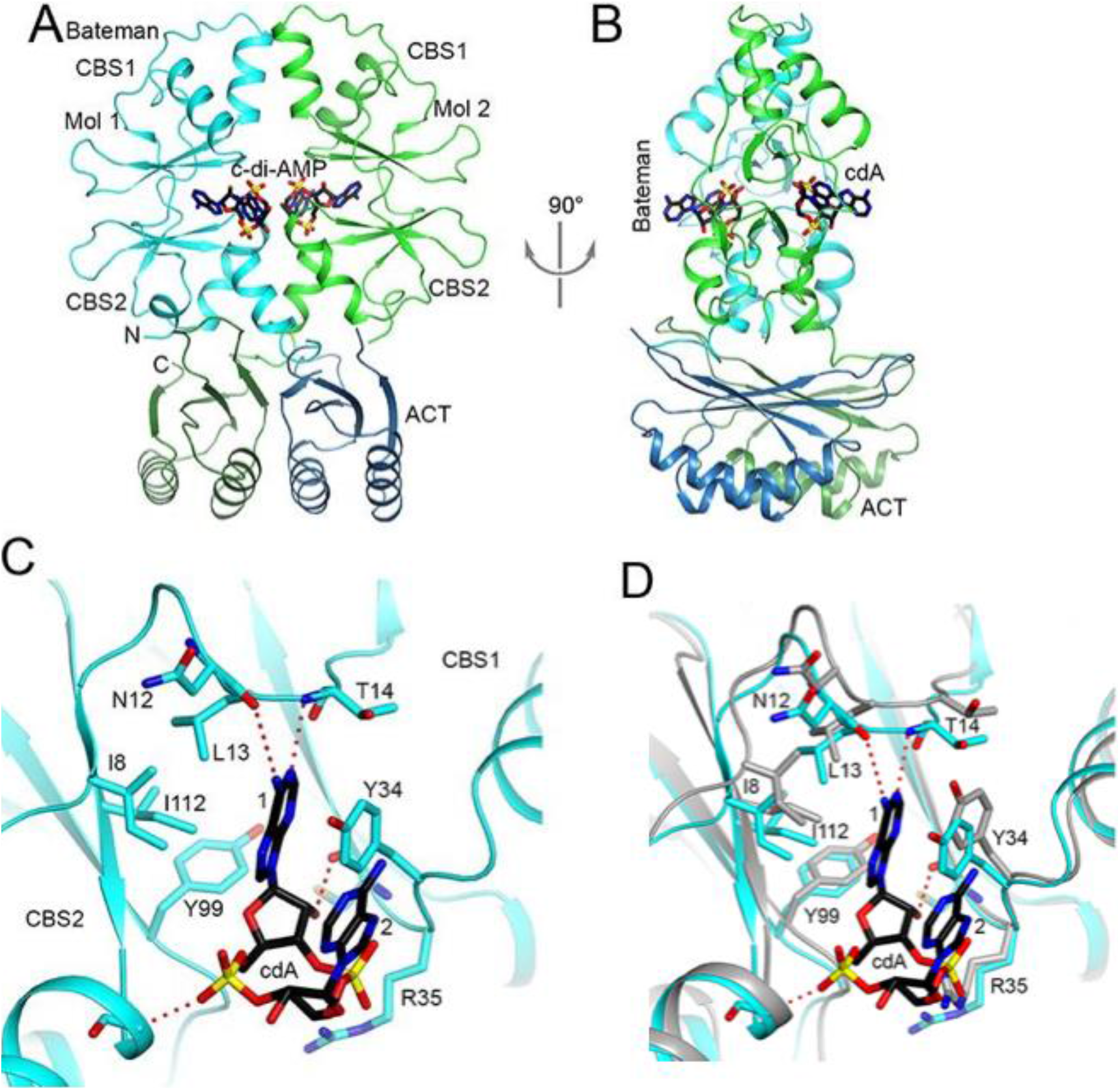
Crystal structure of Lmo0553 in complex with c-di-AMP. **(A)** Schematic drawing of the Lmo0553 dimer bound to two c-di-AMP (cdA) molecules. The CBS motifs and the ACT domain of one monomer are colored in cyan and dark blue, respectively, and those of the other monomer in green and dark green. **(B)** Schematic drawing of the Lmo0553 dimer bound to two c-di-AMP (cdA) molecules, viewed after a 90° rotation around the vertical axis. **(C)** Detailed interactions between Lmo0553 and c-di-AMP. The first and second nucleotides of c-di-AMP are labeled 1 and 2, respectively. Hydrogen-bonding interactions are shown as a dashed line (red). **(D)** Conformational changes in the c-di-AMP binding site. Overlay of the structure of Lmo0553 in complex with c-di-AMP (in color) with that of free Lmo0553 (in gray). All structure figures were produced with PyMOL (www.pymol.org).

### Large conformational changes upon c-di-AMP binding guide biosensor design

Although elucidating the native physiology of Lmo0553 remains a subject of investigation, our detailed understanding of the molecular consequences of c-di-AMP binding led us to hypothesize that Lmo0553 could serve as a scaffold for development of a Förster resonance energy transfer (FRET) biosensor. FRET is exquisitely sensitive to angstrom level changes, therefore we compared the structure of free Lmo0553 to that of the c-di-AMP complex to identify conformational differences (Fig. 2A and 2B). After the Bateman domain of one monomer is overlaid, the orientation of the Bateman domain in the other monomer differs by 13° (Fig. 2B). In addition, the orientation of the ACT domain differs by 12° (Fig. 2A).

**Figure 2:**
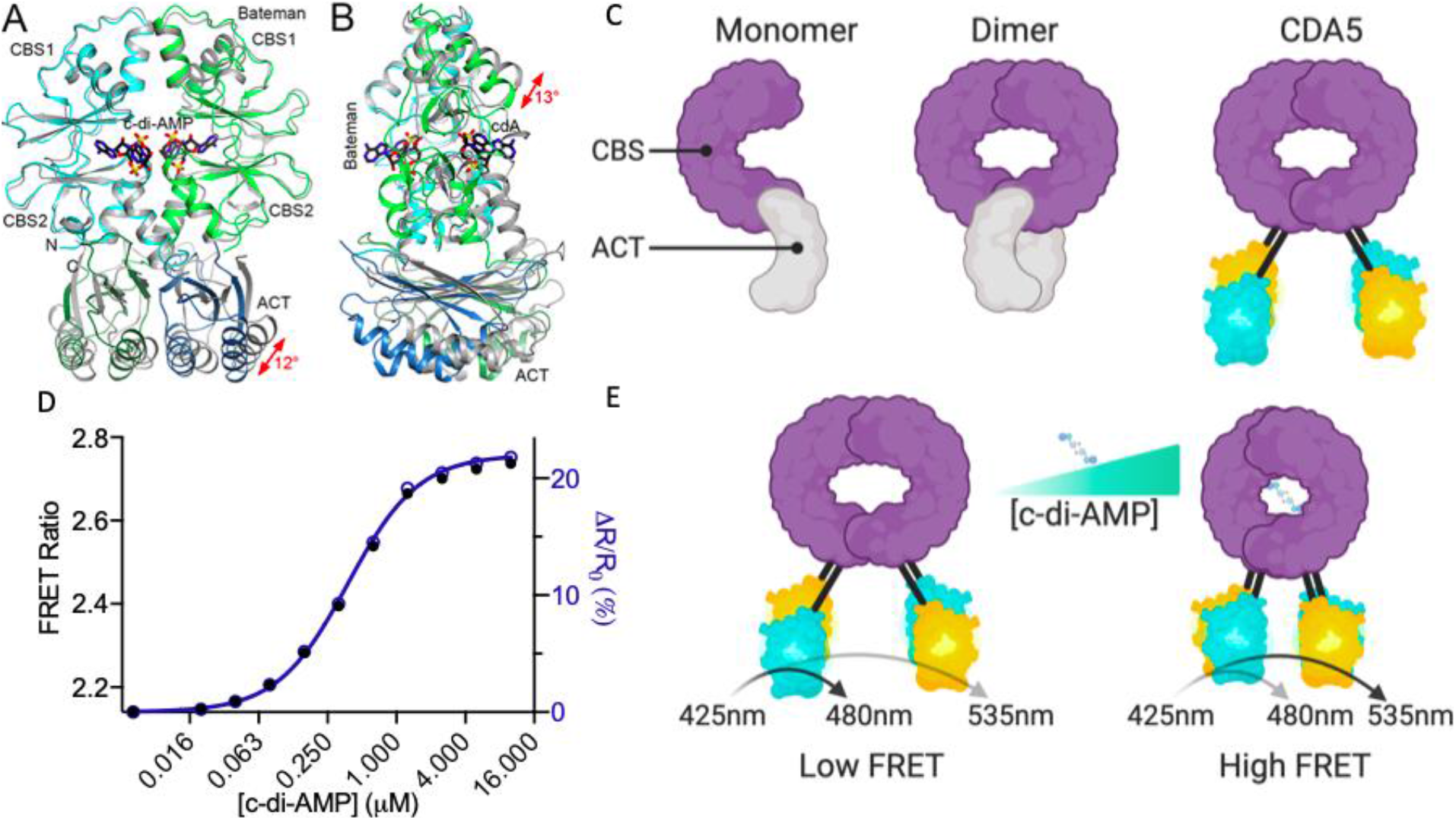
Binding c-di-AMP induces a large structural change in Lmo0553 guiding development of CDA5. **(A)** Overlay of the structure of Lmo0553 in complex with c-di-AMP (in color) with that of free Lmo0553 (in gray). The Bateman domain of one monomer (cyan) is overlaid, in order to visualize the changes in the position of the other domains in the dimer. The conformational change in the ACT domain is indicated by the red arrow. **(B)** Same as panel A, but viewed after a 90° rotation around the vertical axis. The conformation change in the CBS domain is indicated by the red arrow. **(C)** Model of the restructuring used to generate the CDA5 biosensor **(D)** Recombinant CDA5 FRET response to increasing concentrations of c-di-AMP using 425nm excitation and 480nm and 535nm emission wavelengths. Data presented as individual n=1 data points. **(E)** Schematic of CDA5 FRET increase upon c-di-AMP binding. Panels **A** and **B** were produced with PyMOL (www.pymol.org). Panels **C** and **E** were produced using RioRender (www.biorender.com)

To leverage these structural rearrangements, the FRET pair eCFP and eYFP were respectively fused to the N and C termini of full length Lmo0553. This construct was purified and found to be stable but no FRET response was observed upon the addition of c-di-AMP suggesting that fluorophores were too distant or oriented such that structural rearrangements were not detected. Additional engineering may have allowed for the generation of a FRET biosensor utilizing full length Lmo0553, but without a response to optimize, these efforts would have been arduous and potentially unsuccessful.

Reanalyzing the crystal structures, we noted that truncation of the ACT domain might lead to a better biosensor scaffold. The N-terminal Bateman domain of Lmo0553 is distinct from the ACT domain, encompasses the entire binding domain, undergoes large structural changes upon c-di-AMP binding, and, if isolated, would bring fluorophores into close contact increasing the potential to detect FRET ratio changes (Fig. S1B). To generate this second iteration biosensor, Lmo0553 was truncated at N-127 and fused to eCFP and eYFP (Fig. 2C) to best leverage expected large conformational shifts and avoid disruption of the globular domain.

This second iteration c-di-AMP biosensor was recombinantly produced and found to be stable in solution. Excitingly, increasing concentrations of c-di-AMP caused a robust, 23% FRET increase with an EC50 of 0.38 μM (Fig. 2D). Thus, when this biosensor binds c-di-AMP, the chromophores of the flanking fluorophores come in closer contact allowing for increased energy transfer producing an elevated FRET response (Fig. 2E). Satisfied by the magnitude of FRET response, we named this biosensor CDA5 (<u>c</u>yclic-<u>d</u>i-<u>A</u>MP biosensor based on Lmo0<u>5</u>53) and sought to characterize it biochemically.

### Affinity and specificity of CDA5

Although initial FRET response assays suggested that CDA5 retains physiologically relevant biochemical parameters, we sought to further validate this assumption by comparing it directly to full length Lmo0553. DRaCALA analysis [^54^] using [^32^P]-labeled c-di-AMP was employed which revealed an apparent Kd of 4.83 μM and 5.87 μM for Lmo0553 and CDA5 respectively (Fig 3A). This result demonstrates unaltered affinity for c-di-AMP supporting our hypothesis that neither truncation nor fusion with eCFP or eYFP altered ligand binding thermodynamics. To ensure that CDA5 engineering did not alter the previously reported specificity of Lmo0553, the DRaCALA assay was again employed [^13^]. This was done by competing bound radiolabeled c-di-AMP with excess unlabeled nucleotides. CDA5 and Lmo0553 both demonstrated exquisite specificity for c-di-AMP, as only unlabeled c-di-AMP but no other nucleotide in a wide array of monophosphate, triphosphate, cyclic, and dicyclic purine containing nucleotides could compete off radioactive c-di-AMP (Fig 3B, S1B). These data support our hypothesis that CDA5 engineering did not alter critical biochemical parameters.

**Figure 3:**
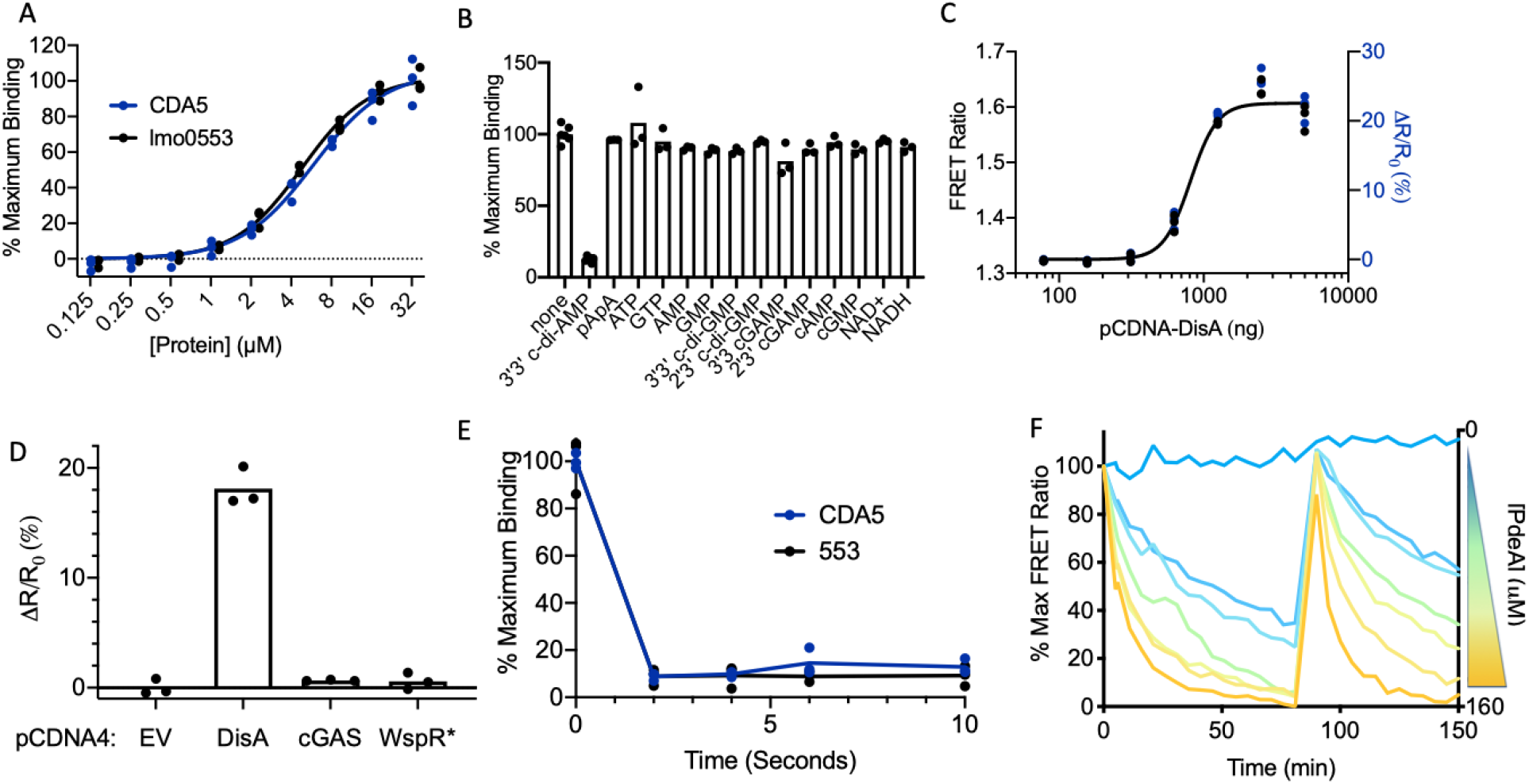
CDA5 retains native and physiologically relevant binding characteristics. **(A)** DRaCALA radioactive nucleotide binding assay of CDA5 (blue) and full length Lmo0553 (black) using ~1nM [^32^P] labeled 3’3’-c-di-AMP. Data fit to a nonlinear curve. Radioactive c-di-AMP bound Lmo0553 and CDA5 at 4.83 μM and 5.87 μM respectively. **(B)** DRaCALA radioactive nucleotide binding assay of CDA5 using ~1nM [^32^P] labeled 3’3’-c-di-AMP in the presence of excess (500 μM) unlabeled nucleotides. Corresponding full length Lmo0553 competition assay is Fig S3B. **(C)** HEK293T cells stably expressing CDA5 were transfected with increasing concentrations of pCDNA4-DisA and analyzed for FRET response by flow cytometry. **(D)** HEK293T cells stably expressing CDA5 were transfected with 2000 ng of expression vectors for DisA, cGAS, WspR*, or empty vector and analyzed for FRET response by flow cytometry. **(E)** DRaCALA radioactive nucleotide binding assay time course of CDA5 (blue) and full length Lmo0553 (black) using ~1nM [^32^P] labeled 3’3’-c-di-AMP in the presence of excess (500 μM) unlabeled c-di-AMP. **(F)** PdeA phosphodiesterase activity assay time course in the presence of decreasing concentrations of recombinant PdeA (2 fold dilutions from 160μM) monitored using CDA5. 2 μM c-di-AMP was re-spiked into the solution at 90 minutes. In panels **A-E** data are presented as n=3 biological replicates. Panel **F** is presented as individual n=1 data points connected by a line.

To further interrogate CDA5 specificity, we utilized a complex cytosolic environment where an array of cyclic dinucleotide (CDN) cyclases could be reliably ectopically expressed. Specifically, we recently engineered a biosensor capable of broadly detecting CDNs, particularly 2’3’-cGAMP, in the eukaryotic cytosol and adapted this system for CDA5 [^44^]. We observed a titratable FRET response to the c-di-AMP cyclase DisA and, as expected, no response to high levels of either WspR*, a 3’3’-c-di-GMP cyclase, or cGAS, a 2’3’-cGAMP cyclase, further validating the specificity of CDA5 for c-di-AMP (Fig 3C and 3D).

### CDA5 Reversibility

To be a reliable reporter, CDA5 must rapidly respond to both increasing and decreasing concentrations of c-di-AMP. CDA5 relies on a native effector protein scaffold which we hypothesized would rapidly respond to nucleotide second messenger fluctuations in order to carry out post-translational responses. We found that FRET responses were stable immediately upon addition of c-di-AMP providing evidence of a k_on_ less than the sampling limit of 10 seconds. Similarly, the k_off_ rate was also unable to be quantitated due to sampling limitations. Utilizing the DRaCALA specificity assay, excess unlabelled c-di-AMP was added to both CDA5 and Lmo0553 pre-bound with [^32^P]-labeled c-di-AMP and immediately sampled for binding. Unlabeled c-di-AMP competed off bound [^32^P]-labeled c-di-AMP completely within two seconds providing evidence of a k_off_ less than this time interval (Fig 3D). Despite not being able to quantify k_on_ and k_off_ rates with these assays, our results suggest that CDA5 can rapidly report on c-di-AMP fluctuations.

To further investigate the ability of CDA5 to repetitively detect c-di-AMP increases and decreases, similar to what we hypothesize is occurring in bacterial cells, we cycled c-di-AMP in solution by increasing the concentration using a bolus of c-di-AMP and decreasing the concentration using the c-di-AMP phosphodiesterase, PdeA. As expected, we observed that: PdeA caused a FRET decrease proportional to enzyme concentration, addition of a bolus of c-di-AMP restored the FRET response, and finally, the FRET response decreased again proportional to the concentration of PdeA (Fig 3E). This assay provides evidence that CDA5 is responsive to dynamic c-di-AMP fluctuations and also that CDA5 is a useful platform for kinetic investigations *in vitro*.

### Development of a c-di-AMP blind control

Characterization of CDA5 indicated that it can reliably quantitate c-di-AMP in simplified systems but, recognizing that complexities can occur in native systems, we sought to develop a point mutant control version of CDA5 that does not bind c-di-AMP. Such a control would provide the capacity to separate FRET signal due to bonafide changes in c-di-AMP from other phenomena such as fluorophore quenching and protein-protein interactions.

Analysis of the crystal structure of Lmo0553 identified tyrosine-34 coordination of c-di-AMP binding via p-stacking (Fig 1C). We hypothesized that by replacing the stabilizing tyrosine ring with an alanine (Y34A), c-di-AMP binding would be minimized in a manner unlikely to disrupt protein stability. Recombinant Y34A CDA5 was found to be stable in solution and analyzed for binding using the DRaCALA binding assay. As hypothesized, c-di-AMP bound to CDA5 but not the Y34A CDA5 point mutant control (Fig 4A). Next, increasing concentrations of c-di-AMP were added to purified CDA5 and Y34A CDA5 and analyzed by plate reader assay for FRET response. CDA5 but not Y34A CDA5 produced a robust FRET increase upon the addition of c-di-AMP (Fig 4B). Together these results suggest that Y34A CDA5 remains unbound and in the apo conformation even in the presence of c-di-AMP.

**Figure 4:**
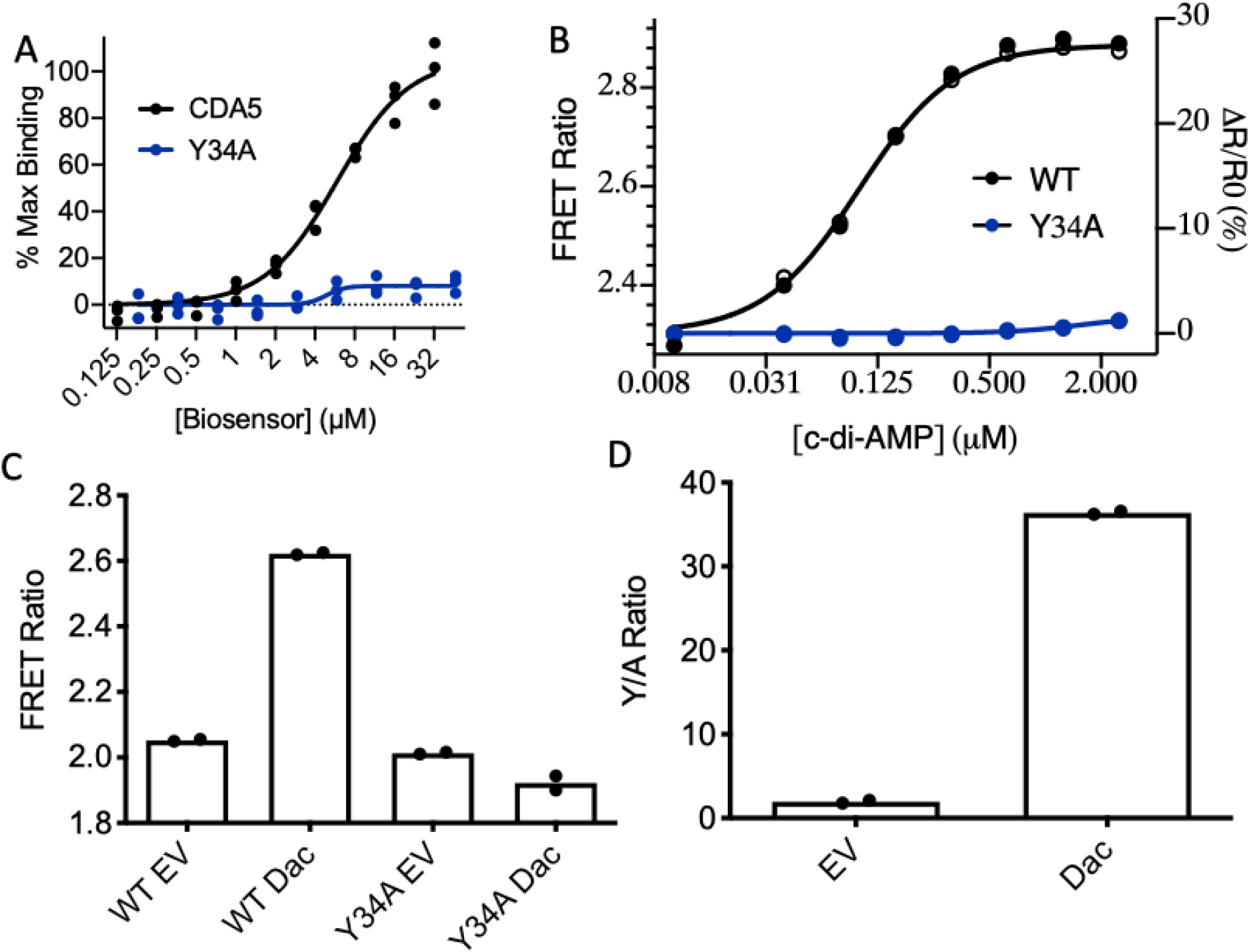
CDA5 Y34A is an important c-di-AMP blind control. **(A)** DRaCALA radioactive nucleotide binding assay of CDA5 WT (black) and Y34A (blue) using ~1nM [^32^P] labeled 3’3’-c-di-AMP. **(B)** Recombinant CDA5 WT (black) and Y34A (blue) FRET response to increasing concentrations of c-di-AMP. **(C)** BL21 (DE3) E. coli transformed with pET15b-CDA5 (WT or Y34A) and pBAV-dacA or empty vector then analyzed by flow cytometry. **(D)** Data in panel C converted into Y/A ratio. Data in panel A is presented as n=3 biological replicates. Panel **B** is presented as individual n=1 data points. Data in panels **C** and **D** are presented as n=2 biological replicates.

We next sought to validate Y34A CDA5 in *Escherichia coli* which is a complex model bacterial organism that does not naturally produce c-di-AMP but can be made to ectopically express a c-di-AMP cyclase and synthesize c-di-AMP. Thus, *E. coli* was transformed with a plasmid to express WT CDA5 or Y34A CDA5 as well as a second plasmid encoding the soluble domain of the c-di-AMP cyclase, DacA or an empty vector and then analyzed for FRET response by flow cytometry (Fig S3A). *E. coli* carrying WT CDA5 produced a robust FRET response while Y34A CDA5 had an, albeit minor, FRET decrease (Fig 4C). We hypothesize that the minor FRET response is due to altered levels of non specific protein-protein interactions. Regardless, these results reinforce the utility of a nonbinding control such as Y34A CDA5 control to provide confidence that observed FRET responses are due to changes in ligand concentration and not other phenomena. It is often useful to also calculate the ratio of WT CDA5 to Y34A CDA5 FRET responses into a ‘Y/A Ratio’ (Fig 4D). This metric, both combines the control biosensor data and declutters data making results more clear.

### CDA5 detects unimodal *Bacillus subtilis* responses to media alteration

CDA5 was then expressed in *Bacillus subtilis* to study native c-di-AMP regulation. *B. subtilis* is a model organism encoding a large array of c-di-AMP cyclases and phosphodiesterases but not a homologue to Lmo0553 which could cause disruptive heterodimerization. We found that rich media like LB is necessary to attain robust CDA5 expression but, at the same time, it is also important to use media with low autofluoresce to clearly quantitate FRET ratios. Due to the necessity of rich media to express CDA5, back dilution into minimal media was not ideal due to the large metabolism changes required to transition to fully biosynthetic growth. We found that we could back dilute *B. subtilis* expressing CDA5 into 10% LB media which retains a complex nutrient content and also has low enough autofluorescence to clearly quantitate FRET by flow cytometry.

In this diluted LB media, we tracked FRET changes overtime and observed a steady FRET decrease during the entire 180 minutes of growth until the WT and Y34A sensor had nearly the same FRET ratio (Fig 5A). To verify this unexpected result, we split our sample to simultaneously detect c-di-AMP using our flow based FRET assay and the gold standard for c-di-AMP quantification, mass-spectrometry. These results were plotted on XY plots graphing c-di-AMP measured by mass spectrometry versus either the WT CDA5 FRET ratio (Fig 5B) or the Y/A ratio combining WT CDA5 and Y34A CDA5 ratios (Fig 5C). Although both the WT CDA5 FRET ratio and the Y/A ratio both correlated with c-di-AMP as measured by mass spec, the Y/A ratio highly correlated with c-di-AMP highlighting the importance of both the Y34A CDA5 control and the Y/A metric.

**Figure 5:**
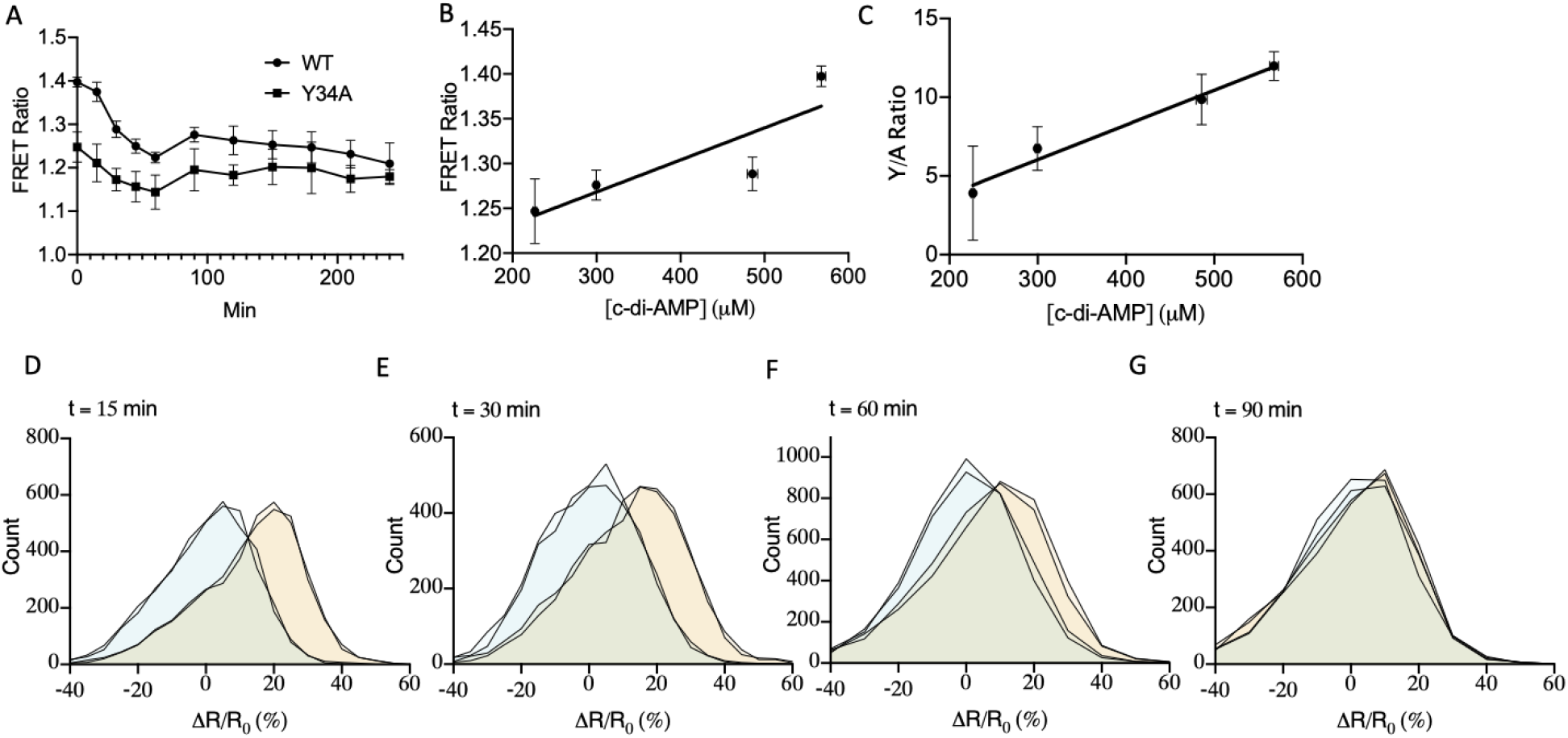
CDA5 allows for the detection of native c-di-AMP dynamics in single cells. **(A)** B. subtilis expressing CDA5 WT (black) and Y34A (blue) back diluted into 10% LB media and grown at 37°C and analyzed for FRET response by flow cytometry **(B)** CDA5 WT FRET ratio in panel A plotted versus c-di-AMP quantitated by mass spectrometry. **(C)** CDA5 Y/A ratio in panel A plotted versus c-di-AMP quantitated by mass spectrometry. **(D-G)** CDA5 WT FRET ratio of single cells divided by the average CDA5 Y34A FRET ratio plotted as histograms at indicated time points. Data in panels **A-C** are presented as mean and standard deviation of n=3 biological replicates. Data in panels **D-G** are presented as histograms consisting of individual data points of n=2 replicates. Blue and orange histograms respectively represent CDA5 Y34A and CDA5 WT.

Flow cytometry allows for the collection of an enormous amount of informative single cell data providing an excellent opportunity to understand more about c-di-AMP dynamics. To understand more about the mechanics of this population level c-di-AMP decrease, WT CDA5 and Y34A CDA5 single cell data was normalized by the control biosensor’s population average and plotted as histograms (Fig 5D-G). This analysis revealed a progressive unimodal population shift as the WT biosensor more closely overlays Y34A CDA5 over time, tracking with population level results. This data also suggests that the vast majority, if not all, of the *B. subtilis* cells are responding in a coordinated fashion to readjust to media conditions.

### CDA5 detects unimodal c-di-AMP differences between *Bacillus subtilis* mutants

To better contextualize the biologic meaning of the FRET ratios detected in WT *B. subtilis*, we next sought to detect c-di-AMP differences between mutants defective in either a c-di-AMP cyclase or phosphodiesterase. As expected, Δ*pgpH*, a mutant lacking the phosphodiesterase PgpH, had higher Y/A ratios due to elevated c-di-AMP; while a mutant lacking the cyclase DisA, Δ*disA*, had lower Y/A ratios due to reduced c-di-AMP production (Fig 6A). All three strains experienced different degrees of FRET ratio decreases. Δ*disA* decreased most quickly, followed by WT, and Δ*pgpH* decreased minimally and remained elevated throughout the experiment. At the single cell level Y/A ratios for all strains again decreased in a unimodal fashion following the population average (Fig 6B-C S6A-F). More experiments will be required to elucidate the mechanism by which Δ*pgpH* but not other strains retain elevated c-di-AMP in reduced LB media. Importantly, this data provides convincing evidence that CDA5 detects physiologically relevant differences in c-di-AMP in *B. subtilis* is a useful tool to investigate bacterial c-di-AMP dynamics.

**Figure 6:**
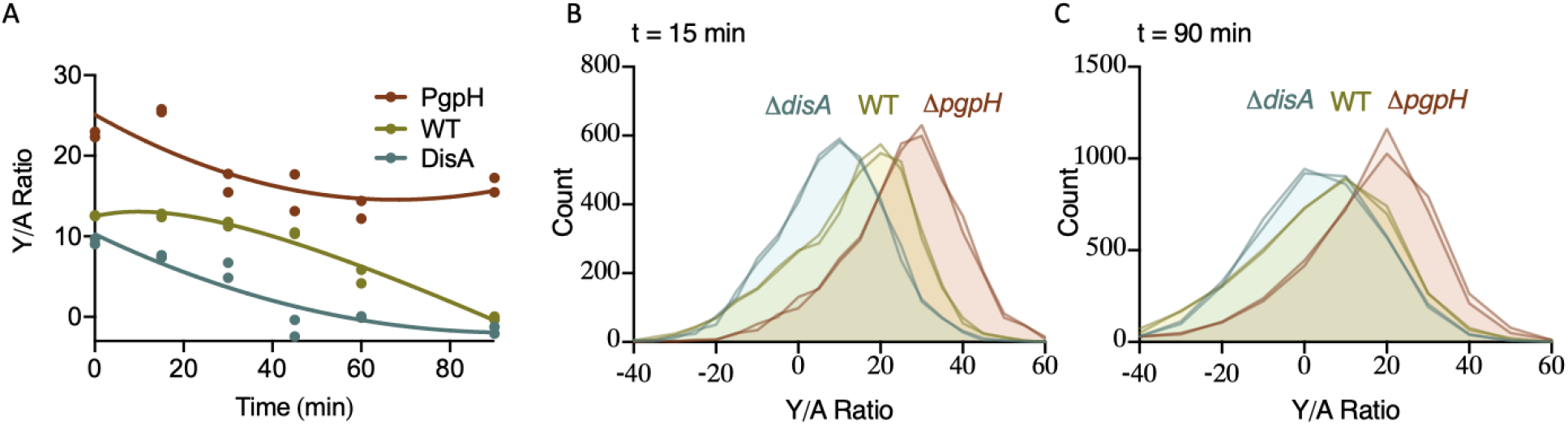
CDA5 detects varied c-di-AMP dynamics in B. subtilis mutant. **(A)** WT (yellow), ΔpgpH (red), and ΔdisA (blue) *B. subtilis* back diluted into 10% LB media and grown at 37°C and analyzed for FRET response by flow cytometry presented as Y/A ratios **(B-C)** CDA5 WT FRET ratio of single cells divided by the average CDA5 Y34A FRET ratio plotted as histograms at indicated time points. Data in all panels are presented as individual data points of n=2 biological replicates.

## DISCUSSION

In this work we describe the development of CDA5, a FRET based biosensor capable of detecting the essential signaling molecule c-di-AMP within individual bacterial cells. Through rational design based on the Lmo0553 crystal structure, we were able to generate a stable biosensor which retains relevant native binding characteristics as well as a nucleotide blind control which improves measurement accuracy. *In vivo*, CDA5 allowed for the detection of c-di-AMP differences over time and between mutants of *B. subtilis*. Interestingly, analysis at the single cell level revealed unimodal c-di-AMP shifts providing evidence that the entire bacterial population responds in a coordinated fashion. This is particularly notable because it is in contrast to the bacterial nucleotide second messenger c-di-GMP which is regulated in a largely bimodal manner[^38–40,42^].

In addition to its ability to monitor native c-di-AMP regulation, CDA5 is also capable of facilitating diverse investigations of c-di-AMP in other contexts. CDA5 is easy to produce, specific, and provides a kinetic readout making it a good platform to study c-di-AMP enzymology. For example, CDA5 can be used to detect protein-protein or small molecule dependent activation or inhibition of c-di-AMP cyclases and phosphodiesterases *in vitro*. Such work may allow for the development of new antibiotics targeting this essential signaling molecule. Some interactions, especially those of membrane associated proteins, are difficult to model with recombinant protein and are better investigated in an ectopic cytosolic environment [^27^]. CDA5 expressed in such a model system would facilitate these investigations. For example, CDA5 expressed in *E. coli* would provide a platform for the interrogation of important protein-protein interactions controlling c-di-AMP synthesis and degradation. Additionally, c-di-AMP is known to be detected by but not produced by eukaryotic cells during infection and perhaps also by certain bacteria like *Pseudomonas aeruginosa* in multicellular environments [^30,55^]. CDA5 expressed by these organisms may be able to detect accumulated c-di-AMP accelerating these interesting investigations.

Although alternative uses are promising, the primary motivation for the development of the CDA5 biosensor was to more thoroughly and efficiently investigate native c-di-AMP regulation. Current tools, including our recently developed CDA-Luc assay, are well suited for detecting c-di-AMP at the population level but the ability to measure c-di-AMP kinetically or at the single cell level is lacking. Thus, CDA5 is a major advance which allows for a wealth of new investigations. A major finding using the c-di-GMP biosensor was mother-daughter cell heterogeneity which allows for different roles within a population [^38,39^]. Similarly we hypothesize that there will be replication phase dependent c-di-AMP fluctuations in bacteria that naturally produce c-di-AMP. However, rather than mother-daughter cell heterogeneity, we hypothesize that the intracellular concentration of c-di-AMP is linked to periods of rapid peptidoglycan synthesis during bacterial elongation due to the close link between c-di-AMP and cell wall homeostasis. Such insights could lead to greater understanding of morphology and virulence differences between c-di-AMP mutants.

Furthermore, the c-di-GMP biosensor was recently utilized to detect regulation of c-di-GMP signaling during macrophage infection [^42^]. Due to the avirulence of mutants which hyper- or hypo- produce c-di-AMP, we hypothesize that similar, if not more profound, regulation is occurring during infection of eukaryotic cells. Such investigations will help elucidate the role of c-di-AMP during infection for a multitude of clinically important organisms including *Streptococcus pneumoniae, Listeria monocytogenes,* and *Mycobacteria tuberculosis* [^13,16,19–21,56^].

CDA5, similarly to previous c-di-GMP biosensor studies [^40^], will also facilitate the identification of diverse environmental stimuli that regulate c-di-AMP at the population and subpopulation level. In this study, we detected a unimodal c-di-AMP response to specific environmental conditions but CDA5 can elucidate a more thorough understanding of c-di-AMP regulation via use of an arrayed media library containing a wide variety of nutrients and stress conditions. In addition to advancing basic biology, such studies may also facilitate the development of clinical interventions which alter intracellular c-di-AMP concentrations.

CDA5 is highly functional in its current form but future development will further improve utility. One such improvement is the use and subsequent optimization of alternative fluorophores. For example, a far-red shifted FRET pair could be utilized allowing for simultaneous non-overlapping fluorescence with BFP-tagged proteins [^52,53,57,58^]. Similarly, a BRET pair could be developed for c-di-AMP detection within mammalian tissue or biofilms [^43,59,60^]. Another type of improvement is modification of the nucleotide binding pocket to increase affinity or alter specificity. For example, it may be possible to engineer additional hydrogen bonding interactions to increase the affinity for c-di-AMP allowing for detection of lower concentrations of nucleotides. Such a sensor would be able to detect low levels of c-di-AMP found within a mammalian cytosol during infection or within a non-c-di-AMP producing bacteria in a biofilm environment. Similarly, the binding pocket of CDA5 may be able to be modified to detect the growing class of CBASS cyclic di-nucleotides produced as part of the anti-bacteriophage response [^61^]. A final type of improvement would be modification of the dimerization domain. It may be possible to reverse polar interactions in the dimerization domain to avoid potentially obfuscating interactions with full length Lmo0553. This would mainly be useful for investigations in *Listeria monocytogenes* or *Enterococcus faecalis* which respectively encode *lmo0553* or a homologue. Alternatively, unless Lmo0553 is the protein of interest, such studies could also be done in a *lmo0553* null background as no phenotype has been identified for this protein to date.

CDA5 is a powerful tool which makes a large class of investigations now feasible. In addition to current capabilities, there are clear next steps to modify CDA5 and apply it to even more diverse studies. Thus, CDA5 holds exceptional promise for accelerating our understanding of the essential bacterial signaling molecule, c-di-AMP.

## METHODS

### Cloning

Primers for CDA5 cloning are listed in Supplementary Table 2, plasmids in Supplementary Table 3, and strains in Supplementary Table 4. CDA5 was generated by subcloning *lmo0553* from the previously generated pET28a-*lmo0553* vector[^13^] using Kapa HiFi polymerase (Kapa Biosystems) using primers 1 and 2. The resulting product was ligated into pET15b-eCFP-12AA-eYFP using Spe1/Kpn1 fast digest restriction endonuclease cloning (Thermo Fisher) and transformed into XL1-Blue chemically competent *E. coli*. To generate pSLIK-CDA5, pET15b-CDA5 was amplified using primers 3 and 4 and added to the BsiW1 (Thermo Fisher) site of pSLIK using InFusion (Takara) then transformed into Stbl3-OneShot competent cells (Thermo Fisher). pET15b-CDA5-Y34A was made using site directed mutagenesis by amplifying the generated pET15b-CDA5 construct using primers 5 and 6 using Kapa HiFi polymerase. PCR product was purified (Promega) and digested using DpnI (NEB) and then transformed into XL1-Blue chemically competent *E. coli*. To generate pHT264-Bs-CDA5, a gBlock codon optimized for *Bacillus subtilis* expression (IDT) was purchased and amplified using primers 7 and 8. The resulting product was ligated into pET15b using Not1 fast digest restriction endonuclease cloning (Thermo Fisher) and transformed into XL1-Blue chemically competent *E. coli* generating pET15b-Bs-CDA5. pET15b-Bs-CDA5-Y34A was generated as above using primers 9 and 10. Finally, pHT264-Bs-CDA5 and pHT264-Bs-CDA5-Y34A were generated by amplifying pET15b-Bs vectors with primers 11 and 12. The resulting product was ligated into pHT264 using BamH1/Smal fast digest restriction endonuclease cloning (Thermo Fisher) and transformed into XL1-Blue chemically competent *E. coli.*

### Protein expression

To obtain *L. monocytogenes* full-length Lmo0553, pET28a-*lmo0553* vector with an N-terminal hexa-histidine tag was transformed into BL21 Star (DE3) cells. The cells were cultured in LB medium with 35 mg/L kanamycin and were induced for 14 h at 20 °C with 1 mM IPTG. A selenomethionine-derivative of Lmo0553 was expressed using a methionine-auxotroph *E. coli* BL21 strain, and the defined medium was supplemented with selenomethionine[^62^]. The protein was purified through nickel-agarose affinity chromatography followed by gel filtration chromatography (S-300, GE Healthcare). The purified protein was concentrated to 30 mg/mL in a buffer containing 20 mM Tris (pH 8.0), 250 mM NaCl, 5% (v/v) glycerol, and 5 mM dithiothreitol, flash-frozen in liquid nitrogen and stored at −80 °C. The N-terminal hexa-histidine tag was not removed for crystallization.

To obtain CDA5, pET15b plasmids encoding CDA5 and CDA5-Y34A were transformed into BL21 Star (DE3) cells. An overnight culture of the transformed bacteria was inoculated into 1 L of LB broth and grown and grown at 37°C until an OD_600_ between 0.5-0.7 at which point protein expression was induced by the addition of 0.1 mM isopropyl β-D-1-thiogalactopyranoside (IPTG) for 16-20 hours at 18°C. The protein was purified using nickel-agarose affinity chromatography (Thermo Scientific). The protein was subsequently buffer exchanged (Cytiva) into Buffer A (40 mM Tris pH 7.5, 100 mM NaCl, 20 mM MgCl_2_, 1mM DTT). Protein samples were tested for purity by SDS-PAGE followed by Coomassie Brilliant Blue staining. Samples were then flash-frozen in liquid nitrogen and stored at −80°C until use in biochemical assays. DisA, RECON, and PdeA were purified the same as above with the exception that they were induced with 1 mM IPTG at 37°C for 4 hours.

### Protein crystallization

Crystals of Lmo0553 in complex with c-di-AMP were grown by the sitting-drop vapor diffusion method at 20 °C. The protein at 15 mg/mL was first incubated with 2.5 mM c-di-AMP for 30 min, and then mixed with reservoir solution containing 23% (w/v) PEG3350, and 0.2 M calcium acetate. The crystals were cryo-protected with the reservoir solution supplemented by 12% (v/v) glycerol and flash-frozen in liquid nitrogen for data collection at 100 K. Crystals of free Lmo0553 were grown by the sitting-drop vapor diffusion method at 20 °C. The protein at 16 mg/mL was mixed with a reservoir solution containing 1.4 M ammonium sulfate, and 0.1 M sodium citrate (pH 5). The crystals were cryo-protected with 20% (v/v) glycerol and flash-frozen in liquid nitrogen for data collection at 100 K.

### Data collection, structure determination and refinement

X-ray diffraction data for Lmo0553 were collected on the X29A beamline at the National Synchrotron Light Source. The diffraction images were processed using HKL2000[^63^]. The structure was solved using the single-wavelength anomalous dispersion (SAD) method with selenomethionine-derivatized crystals, using the program PHENIX[^64^]. Manual rebuilding was carried out in Coot[^65^] and refinement was done with the program Refmac[^66^]. Data collection and refinement statistics are summarized in Table 1.

### Radioactive nucleotide binding assays

[^32^P] 3’3’-cyclic di-AMP was synthesized identically as previously described[^44^]. This nucleotide was then used to perform DRaCALA assays ^54^. Briefly, binding assays were performed in Buffer A at room temperature. To determine binding affinities, two-fold serial dilutions of proteins were incubated with ~1 nM of radioactive 3’3’-cyclic di-AMP for 10 minutes then blotted onto nitrocellulose membranes and allowed to air dry. To determine binding specificity, samples were pre-incubated with 500 μM excess unlabeled nucleotides for 10 minutes followed by incubation with ~1 nM of radioactive 3’3’-cyclic-di-AMP for 10 minutes then blotted onto nitrocellulose membranes and allowed to dry. Finally, to determine the time frame of competition, ~1 nM of radioactive 3’3’-cyclic di-AMP was preincubated with protein for 10 minutes at which point 500 μM of unlabeled 3’3’-cyclic-di-AMP was added, mixed, and blotted onto nitrocellulose membranes every two seconds and allowed to dry. [^32^P] radioactivity was visualized by exposure onto Phosphor-Imager screens, which were developed using a Typhoon FLA 9000 biomolecular imager (GE Healthcare).

### *In vitro* FRET measurements

In all assays, 2 μM of CDA5 or CDA5-Y34A was incubated in black flat bottom half volume opaque 96-well plates (Greiner Bio-One). For nucleotide response assays, two fold dilutions of 3’3’-c-di-AMP (Invitrogen) were made in molecular grade water and added to the protein solution. eCFP and FRET fluorescence was monitored at room temperature using a fluorimeter (BioTek Synergy H1 Hybrid Reader, Biotek Instruments) at 425 nm excitation and read at 480nm and 535nm emission wavelengths for eCFP and FRET respectively. PdeA enzyme activity assay was performed as above with the exception that: Buffer A was supplemented 20 mM MnCl_2_, two fold dilutions of PdeA rather than c-di-AMP were added to each well, samples were spiked with 2 μM c-di-AMP at t=0 and t=90 minutes, and the assay was monitored every 5 minutes.

### Eukaryotic CDA5 specificity assays

Assays were performed similarly as previously described[^44^]. Briefly, Human Embryonic Kidney (HEK) 293T cells were grown in Glutamax Dulbecco’s Modified Eagle Medium (DMEM) (Gibco) supplemented with 10% (v/v) heat-inactivated FBS (HyClone) and 100 U mL^−1^ penicillin and 100 μg mL^−1^ streptomycin and maintained at 37°C and 5% CO2 in a humidified incubator. Self-inactivating lentivirus was made via transfection of a semi-confluent 10 cm dish of HEK293T cells with 4 μg of psPAX2, 2 μg of pCMV-VSV-G, and 4 μg of pSLIK lentiviral vector using Poly(ethyleneimine) (PEI). Growth media was replaced 24 hours after transfection and supernatants collected at 48 and 72 hours, pooled, and filtered through a 0.45 μm filter. Lentivirus was then concentrated with a Lenti-X concentrator (Takara) and added to 4 million HEK293T cells seeded on a 10 cm plate and spinfected for 1 hour at 500X g. After a 24 hour recovery period, media was replaced and supplemented with 2 μg mL^−1^ puromycin (Gibco).

Transduced cells were continually passaged and maintained in selection media containing puromycin. For FRET measurements, 750,000 HEK293T cells transduced with the doxycycline inducible pSLIK-CDA5 plasmid were plated in a 6-well culture dish. The subsequent day, cells were transfected with indicated concentrations of cyclase-encoding plasmids using PEI transfection reagent. One hour after transfection, biosensor expression was induced by the addition of Doxycycline Hydrochloride (Sigma-Aldrich) at 2 μg mL^−1^. The next day, cells were harvested via resuspension in room temperature PBS and CDA5 FRET measurements were collected by FACS analysis. Cells were analyzed using a LSR II flow cytometer (BD) with the following voltages: FSC-350 SSC-240 V500-350 Pacific Blue-420 GFP-400. Data was analyzed using FlowJo software (Tree Star)

### Bacterial FRET measurements

E. coli FRET measurements were obtained by transforming BL21 Star (DE3) cells with pET15b-CDA5 and pET15b-CDA5-Y34A in combination with either pBAVE-EV or pBAVE-DacA and plated on LB Carb 100 μg mL^−1^ Kan 50 μg mL^−1^ plates. An overnight culture of the transformed bacteria was inoculated into 1 L of LB broth and grown and grown at 37°C until an OD_600_ between 0.5-0.7 at which point protein expression was induced by the addition of 0.1 mM isopropyl β-D-1-thiogalactopyranoside (IPTG) for 16-20 hours at 18°C. At this point, cells were spun down and resuspended in room temperature PBS and CDA5 FRET measurements collected by FACS analysis. Cells were analyzed using a LSR II flow cytometer (BD) with the following voltages: FSC-400 SSC-200 V500-500 Pacific Blue-600 GFP-400. Data was analyzed using FlowJo software (Tree Star) B. subtilis FRET measurements were obtained by transforming B. subtilis with pET264-CDA5 and pET264-CDA5-Y34A and plating cells on LB Carb 100 μg mL^−1^ plates and incubated at 37℃. Single colonies were then struck onto LB Carb 100 μg mL^−1^ IPTG 1 mM plates and incubated overnight at 30°C. The resulting single colonies were harvested in 10% LB media (FRET detectable up to 30% LB media) and clumps dissociated by passing cells 2-3 times through a 27 gauge needle. Cells were then grown shaking at 37°C until desired time points at which CDA5 FRET measurements were collected by FACS analysis. Cells were analyzed using a LSR II flow cytometer (BD) with the following voltages: FSC-400 SSC-200 V500-450 Pacific Blue-550 GFP-375. Data was analyzed using FlowJo software (Tree Star).

### Mass spectrometry

The OD_600_ of *B. subtilis* samples was taken. Then, half of the sample was analyzed by FACS analysis and the other half pelleted and frozen for c-di-AMP extraction. Cell pellets were resuspended in 50 μL of 0.5 μM heavy-labeled (C13 N15) c-di-AMP, then mixed with 500 μL of methanol and sonicated. The sample was pooled, centrifuged, and supernatant saved. The resulting pellet was resuspended in 50 μL water then mixed with 500 μL methanol and sonicated again. The solution was centrifuged and the second supernatant pooled with the first. The extract was then dried using a speed vacuum concentrator. The resulting film was resuspended in 50 μL of molecular grade water and measured by mass spectrometry as described[^16^].

### DATA AVAILABILITY

The datasets generated and/or analyzed during the current study are available from the corresponding author on reasonable request.

## ACKNOWLEDGEMENTS

We thank all of the members of the Woodard lab for constructive comments throughout the course of this work. Graphics used in this work were generated using BioRender. A.J.P was supported in part by a Public Health Service, National Research Service Award, T32GM007270, from the National Institute of General Medical Sciences. Research in the Woodward lab was supported by National Institutes of Health Grants 5R01AI139071-02, 1R21AI137758-01 and 1R21AI153820-01. Research in the Tong lab was supported by National Institutes of Health grant AI116669 (to J.J.W. and L.T.).

## AUTHOR CONTRIBUTIONS

A.J.P. and J.J.W. conceived the study. A.J.P., P.H.C., and S.A.Z. designed and performed experiments and analyzed data. A.J.P., P.H.C., L.T., and J.J.W. wrote the manuscript. All authors edited the manuscript.

## COMPETING INTERESTS

The authors declare no competing financial interests.

---

**Figure S1:**
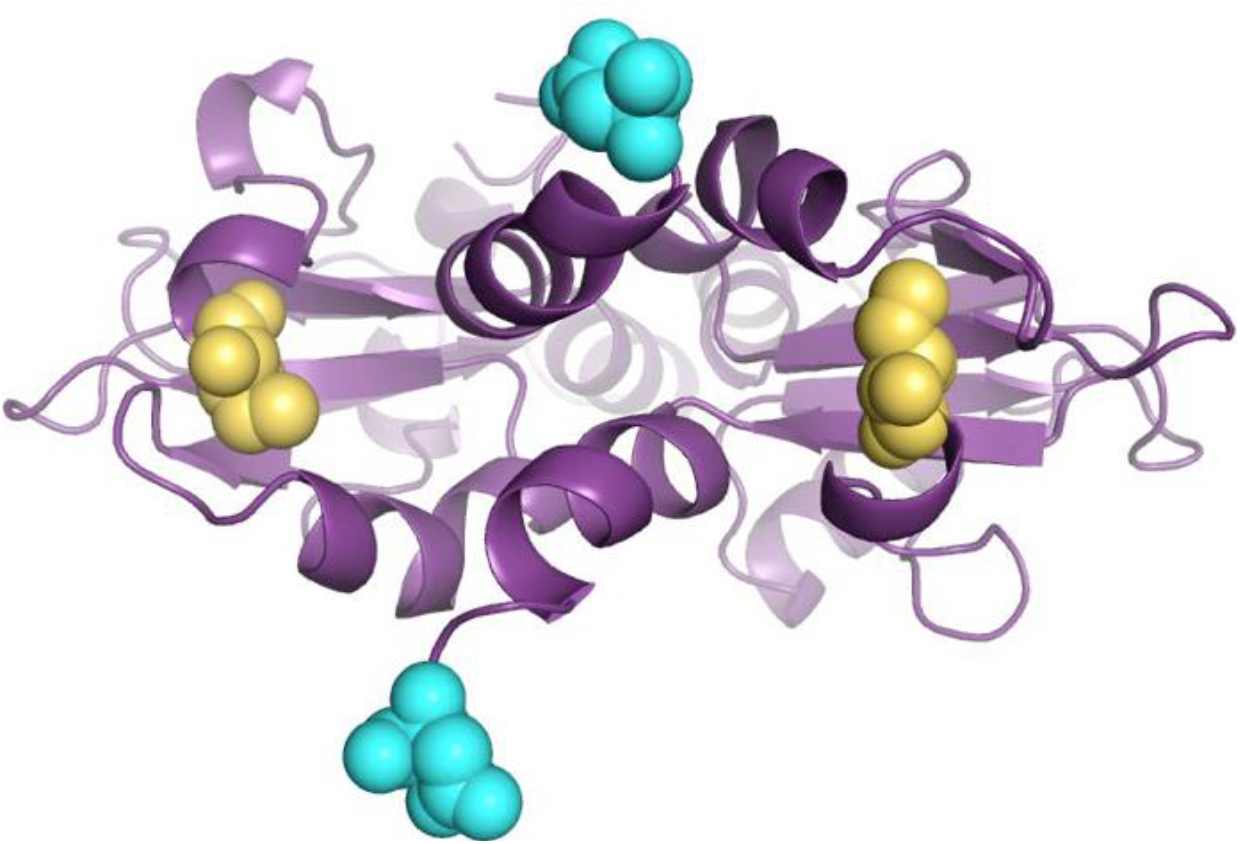
CBS domain structural rearrangement upon c-di-AMP binding Movie models the rearrangement of Lmo0553 upon c-di-AMP binding. Yellow and Cyan denote location of eYFP and eCFP fusion respectively. Made using PyMol (www.PyMol.com)

**Supplemental Figure 2:**
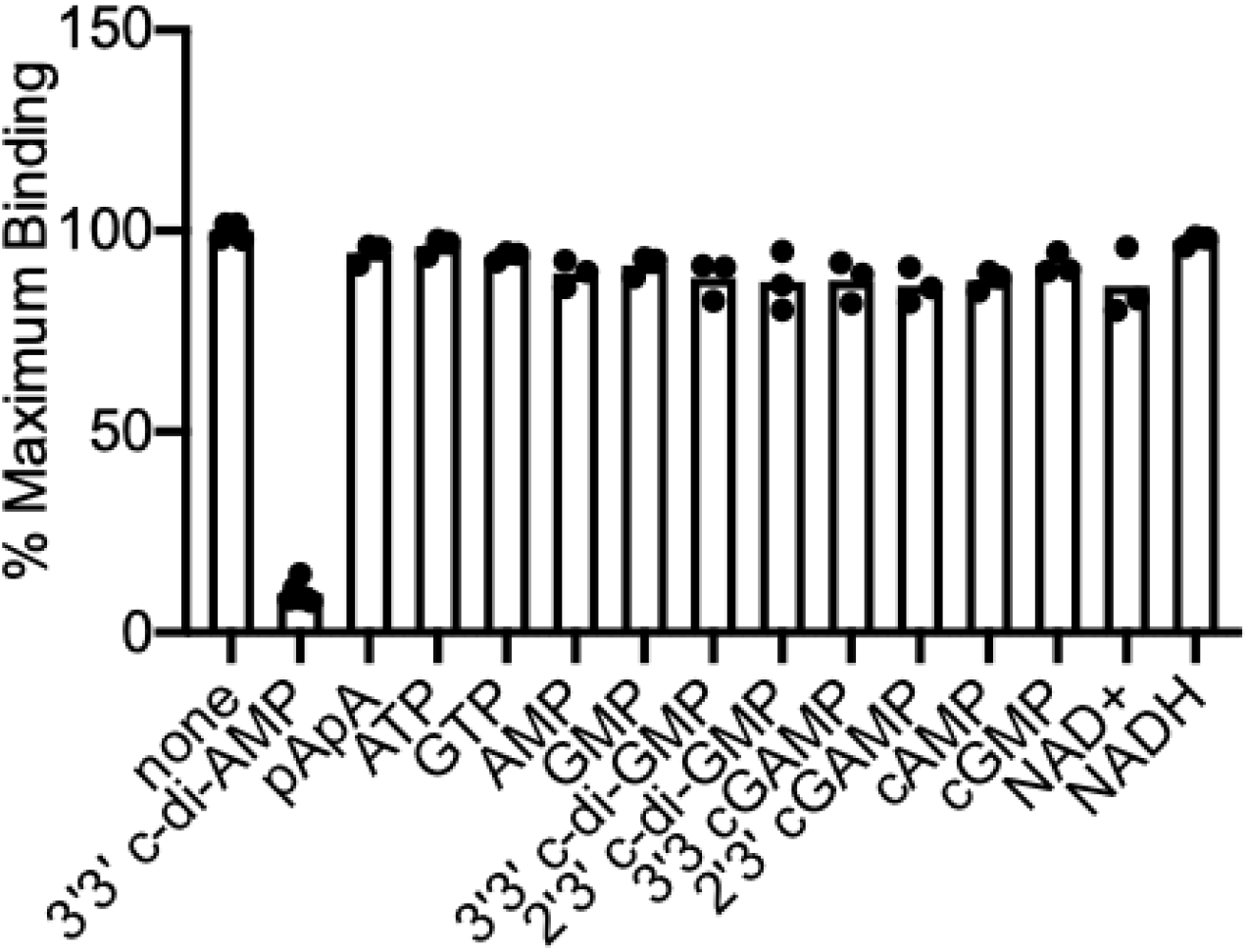
Lmo0553 DRaCALA specificity assay. DRaCALA radioactive nucleotide binding assay of full length Lmo0553 using ~1nM [^32^P] labeled 3’3’-c-di-AMP in the presence of excess (500 μM) unlabeled nucleotides. Data are presented as n=3 biological replicates.

**Supplemental Figure 3:**
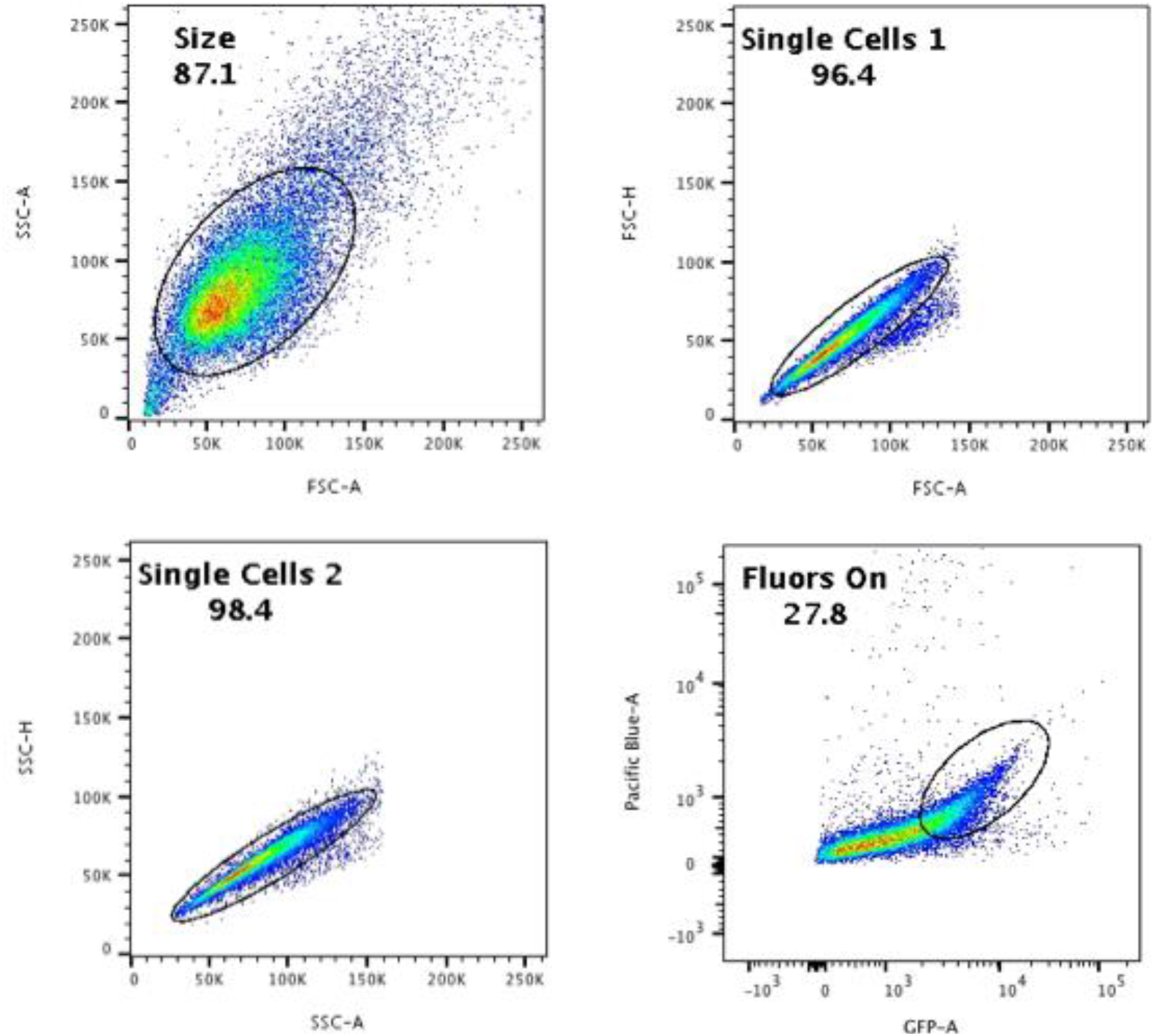
Flow sort method for HEK 293T cells. Cells were sorted using the following gating strategy. All cells in the ‘Fluors On’ gate were analyzed.

**Supplemental Figure 4:**
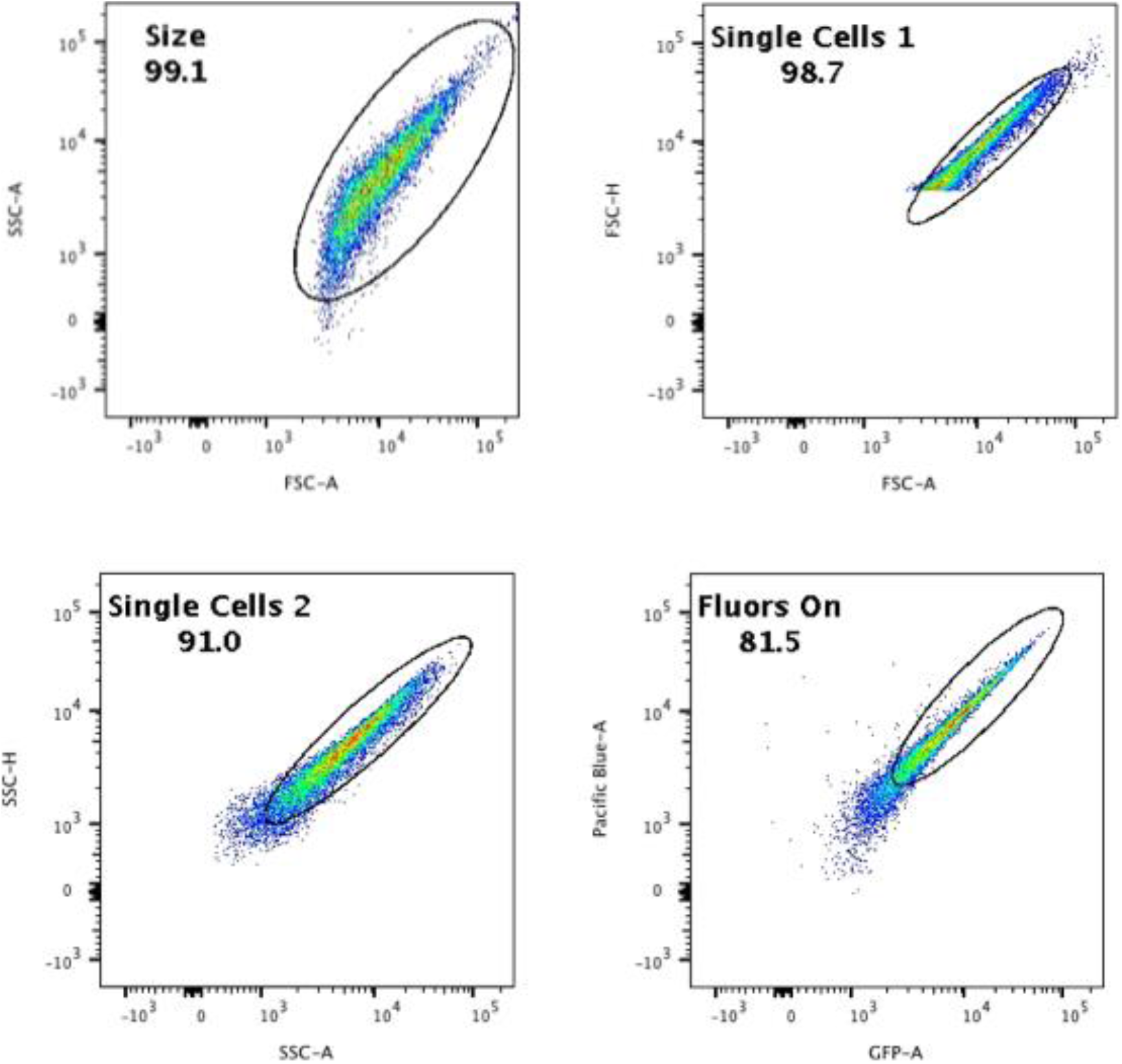
Flow sort method for *E. coli*. Cells were sorted using the following gating strategy. All cells in the ‘Fluors On’ gate were analyzed.

**Supplemental Figure 5:**
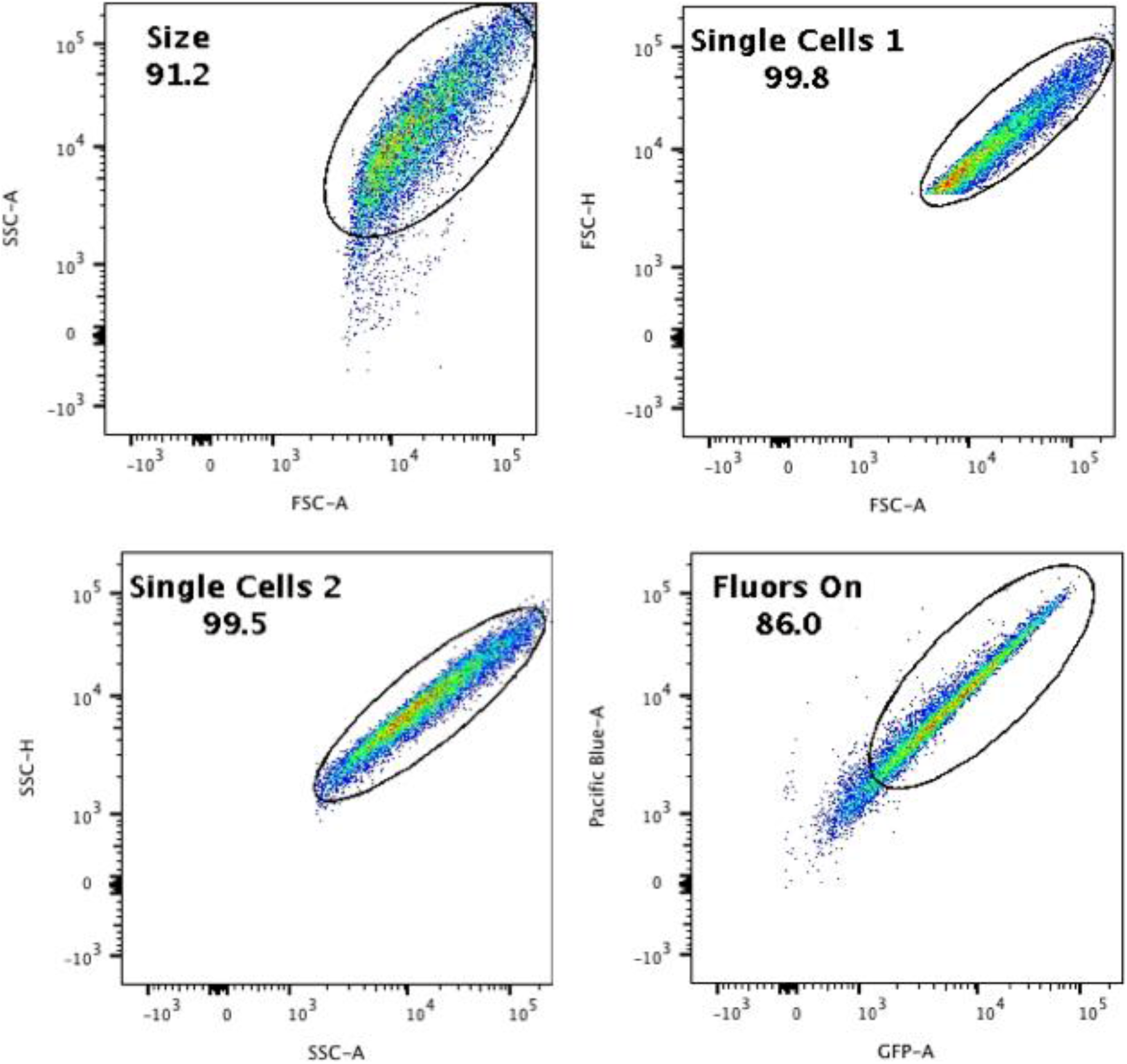
Flow sort method for *B. subtilis*. Cells were sorted using the following gating strategy. All cells in the ‘Fluors On’ gate were analyzed.

**Supplemental Figure 6:**
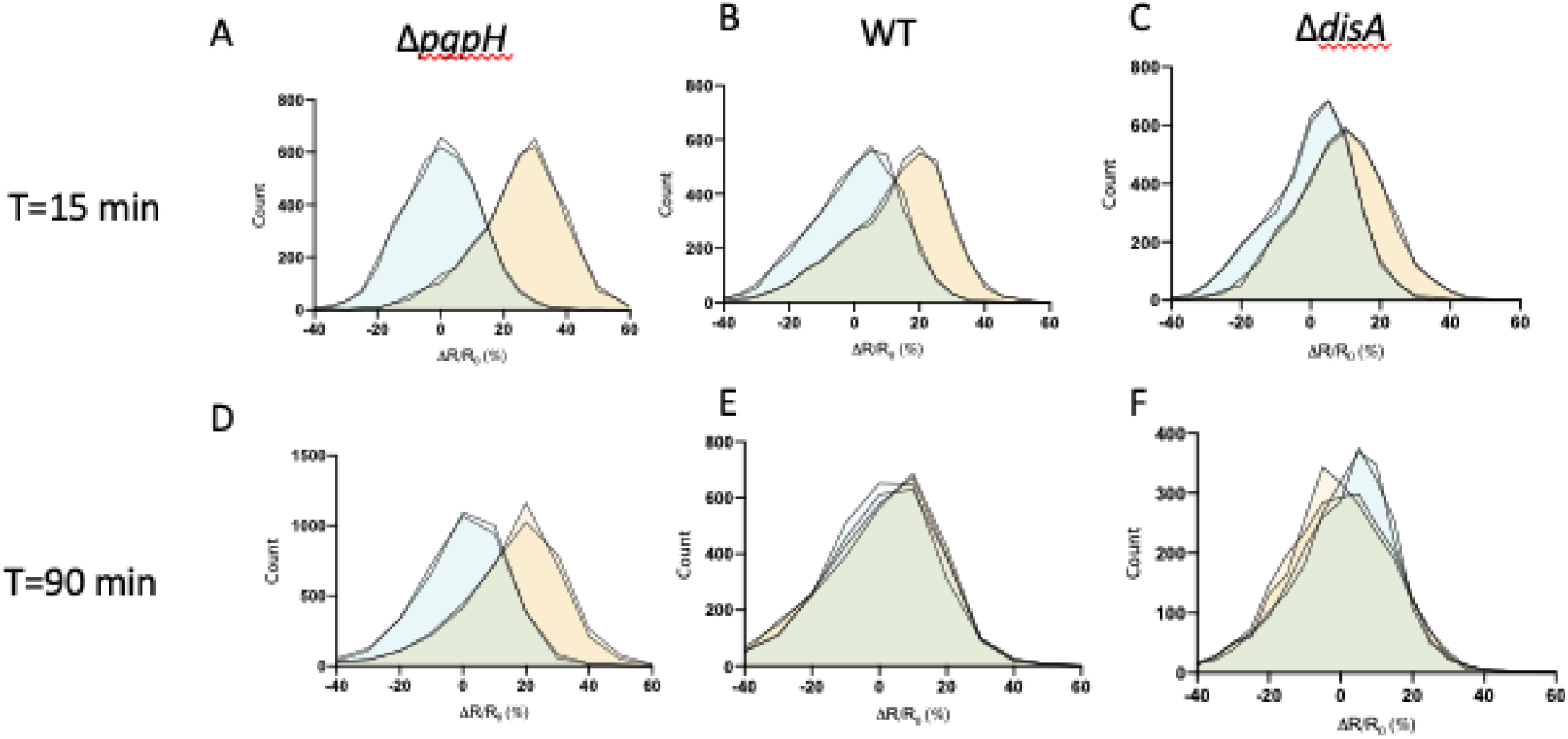
CDA5 WT and CDA5 Y34A single cell data. **(A-F)** CDA5 WT and CDA5 Y34A FRET ratios of single cells divided by the average CDA5 Y34A FRET ratio plotted as histograms of indicated time points and *B. subtilis* strains

